# Systematic multi-trait AAV capsid engineering for efficient gene delivery

**DOI:** 10.1101/2022.12.22.521680

**Authors:** Fatma-Elzahraa Eid, Albert T. Chen, Ken Y. Chan, Qin Huang, Qingxia Zheng, Isabelle G. Tobey, Simon Pacouret, Pamela P. Brauer, Casey Keyes, Megan Powell, Jencilin Johnston, Binhui Zhao, Kasper Lage, Alice F. Tarantal, Yujia A. Chan, Benjamin E. Deverman

## Abstract

Broadening gene therapy applications requires manufacturable vectors that efficiently transduce target cells in humans and preclinical models. Conventional selections of adeno-associated virus (AAV) capsid libraries are inefficient at searching the vast sequence space for the small fraction of vectors possessing multiple traits essential for clinical translation. Here, we present Fit4Function, a generalizable machine learning (ML) approach for systematically engineering multi-trait AAV capsids. By leveraging a capsid library that evenly samples the manufacturable sequence space, reproducible screening data are generated to train accurate sequence-to-function models. Combining six models, we designed a multi-trait (liver-targeted, manufacturable) capsid library and validated 89% of library variants on all six predetermined criteria. Furthermore, the models, trained only on mouse *in vivo* and human *in vitro* Fit4Function data, accurately predicted AAV capsid variant biodistribution in macaque. Top candidates exhibited high production yields, efficient murine liver transduction, up to 1000-fold greater human hepatocyte transduction, and increased enrichment, relative to AAV9, in a screen for liver transduction in macaques. The Fit4Function strategy ultimately makes it possible to predict cross-species traits of peptide-modified AAV capsids and is a critical step toward assembling an ML atlas that predicts AAV capsid performance across dozens of traits.

## Introduction

Engineering novel functions into proteins while retaining desired traits is a key challenge for developers of viral vectors, antibodies, and inhibitors of medical and industrial value ^1–3^. For instance, to be harnessed as a viable gene therapy vector, an adeno-associated virus (AAV) capsid should simultaneously exhibit high production yield and efficiently target the cell type(s) relevant to a specific disease across preclinical models to patients. A common approach for developing AAV capsids with novel tropisms is to funnel a random library of peptide-modified capsids through multiple rounds of selection to identify a few top-performing candidates. This approach has produced modified capsids that more efficiently transduce cells throughout the central nervous system (CNS) ^1,4–9^, photoreceptors ^3^, brain endothelial cells ^10,11^, and skeletal muscle ^12,13^. These rare capsids can then be diversified to screen for even more enhanced tropisms ^4^, high production yield ^4^ or cross-species functionality ^13^. However, variants optimized for one trait may be difficult to co-optimize for other traits, and the protein sequence space is too vast to effectively sample by chance for rare variants that are enhanced across multiple traits. As a result, AAV engineering teams often devote many years and significant resources to developing capsids that ultimately fail to be optimized across multiple traits essential for preclinical and clinical translation.

To identify novel and diverse AAV capsids that simultaneously possess multiple traits relevant to gene delivery (e.g., manufacturability, targeting to disease-relevant cells across host species, detargeting from other cell types), vast capsid sequence spaces must be subjected to systematic and unbiased searches. This quickly becomes intractable using traditional methods as the capsid sequence is increasingly modified. For instance, merely inserting a string of 7 amino acids (a 7-mer) into an AAV9 capsid generates a theoretical sequence space of 1.28 billion variants; inserting a 10-mer instead of a 7-mer extends that space to 10 trillion variants.

To address this challenge, we sought to develop a generalizable ML-guided approach to systematically and simultaneously map 7-mer-modified AAV9 capsid sequences to multiple traits of interest. To generate high-quality data that would enable the training of accurate ML models, it was necessary to first create a low-bias, high-diversity library composed only of capsid variants with high production fitness (Fig. 1). This “Fit4Function” library was subjected to *in vivo* and *in vitro* screens for traits relevant to gene therapy, which, as anticipated, resulted in highly reproducible data that could be used to train robust ML models. The models trained using the Fit4Function data were of sufficient accuracy that they could be leveraged in combination to search the much larger, untested, theoretical high production fitness sequence space *in silico* for rare multi-trait variants. We first demonstrated that six models relating to liver-targeting *in vitro* and *in vivo* could be used in combination to predict sequences that met filters set across all six models. The resulting library of variants exhibited a high 88.5% validation rate, i.e., 88.5% of its variants were experimentally determined to fulfill all six criteria. Despite being trained only on mouse *in vivo* and human *in vitro* data, this combination of six models translated to the macaque. Variants nominated for individual validation performed well across human cells and mice compared to AAV9. These same variants provided more efficient transduction of the macaque liver relative to AAV9 when tested in a pooled library format. Notably, the combination of *in vivo* and *in vitro* functional predictors boosted the precision of cross-species prediction compared to the use of any individual model. In other words, we observed value in training models on data from human cell *in vitro* functional assays to predict variants that exhibit the trait of interest in mice and macaque *in vivo*. The Fit4Function approach allowed us to systematically and readily identify the combination of traits that is most critical in predicting a given function of interest; appropriate screening models can be identified and used to enrich for multi-trait capsids prior to studies in NHPs or clinical trials. This novel strategy can inform intelligent searches for AAV capsids that are performant across species and more likely to translate from preclinical models to investigational human gene therapies.

**Fig. 1.**
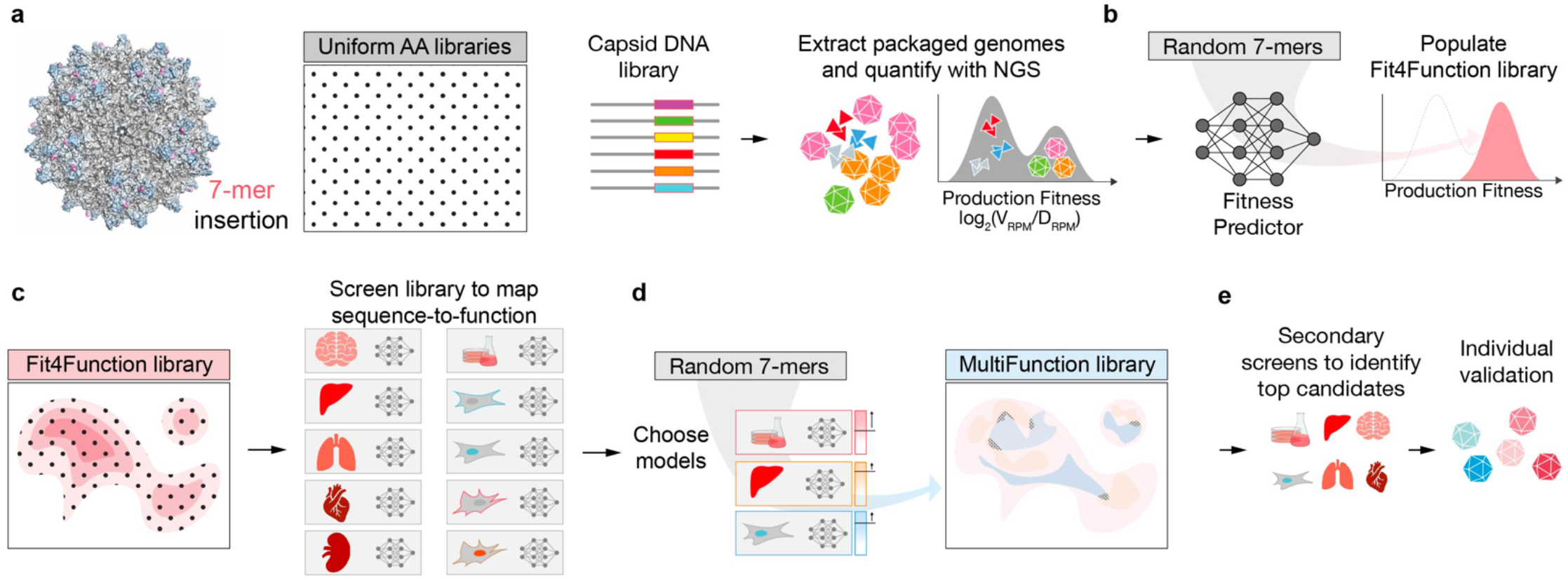
Systematic multi-trait protein optimization paradigm. **(a)**An insertion-modified AAV library that uniformly samples the 7-mer sequence space (1.28 billion possible variants) is designed and used to produce AAV particles. Variant production fitness is measured via NGS of nuclease-resistant *Cap*-containing genomes (V_RPM_) relative to the number of genomes in the DNA library (D_RPM_). **(b)**The production fitness data is used to train a sequence-to-production-fitness ML model that is then used to design the Fit4Function library, which uniformly and exclusively samples the high production fitness sequence space. **(c)**The Fit4Function library can be screened *in vitro* or *in vivo* for functions of interest, and the data are used to derive ML models that predict these functions from random 7-mer sequences. **(d)**The production fitness and functional models are used in combination to populate MultiFunction libraries consisting of variants predicted to perform well across the desired traits (see checkered areas that represent the overlap between the functional sequence spaces of interest). **(e)**The MultiFunction virus libraries are produced and screened for all functions of interest. The top performing variants are then individually validated.

## Results

### High production fitness space mapping

We and others have successfully derived enhanced gene delivery vectors from AAV9 capsids modified through the insertion of 7 amino acids (7-mer) between VP1 residues 588–589. To create an accurate and generalizable sequence-to-production fitness ML model, synthetic modeling and assessment libraries were designed to each consist of 74.5K variants that evenly sample the sequence space (each amino acid was sampled with an equal probability at each position); 10K of the 74.5K are common to both libraries to assess reproducibility across libraries. This is distinct from conventional NNN or NNK (where N is any base and K is a G or T) libraries where millions of variants are synthesized stochastically by uniformly sampling the nucleotide space, which biases toward AAs represented by more codons. Both modeling and assessment libraries were also designed to assess whether codon usage impacts production fitness; each variant is represented by two maximally different nucleotide sequences (7-mer amino acid replicates). We produced both libraries in triplicate, in two separate runs, by two different researchers, for a total of 12 replicates each. The reproducibility (measured by the agreement between replicates) of variant production fitness scores between preparations by different researchers improved as technical and biological replicates were aggregated (Supplementary Fig. 1). Therefore, we performed all subsequent analyses on production fitness scores aggregated across all replicates for each library.

We first assessed whether codon usage impacts the production fitness of identical amino acid variants. If so, it would be necessary to train on the nucleotide sequence space (61^7^ for NNN, 31^7^ for NNK), which is much larger than the amino acid sequence space (20^7^). We observed a high correlation between the fitness scores of 7- mer amino acid replicates in the modeling library (Fig. 2a and Supplementary Fig. 2), suggesting no significant codon usage bias. Therefore, we averaged production fitness across 7-mer replicates for all downstream modeling.

**Fig. 2.**
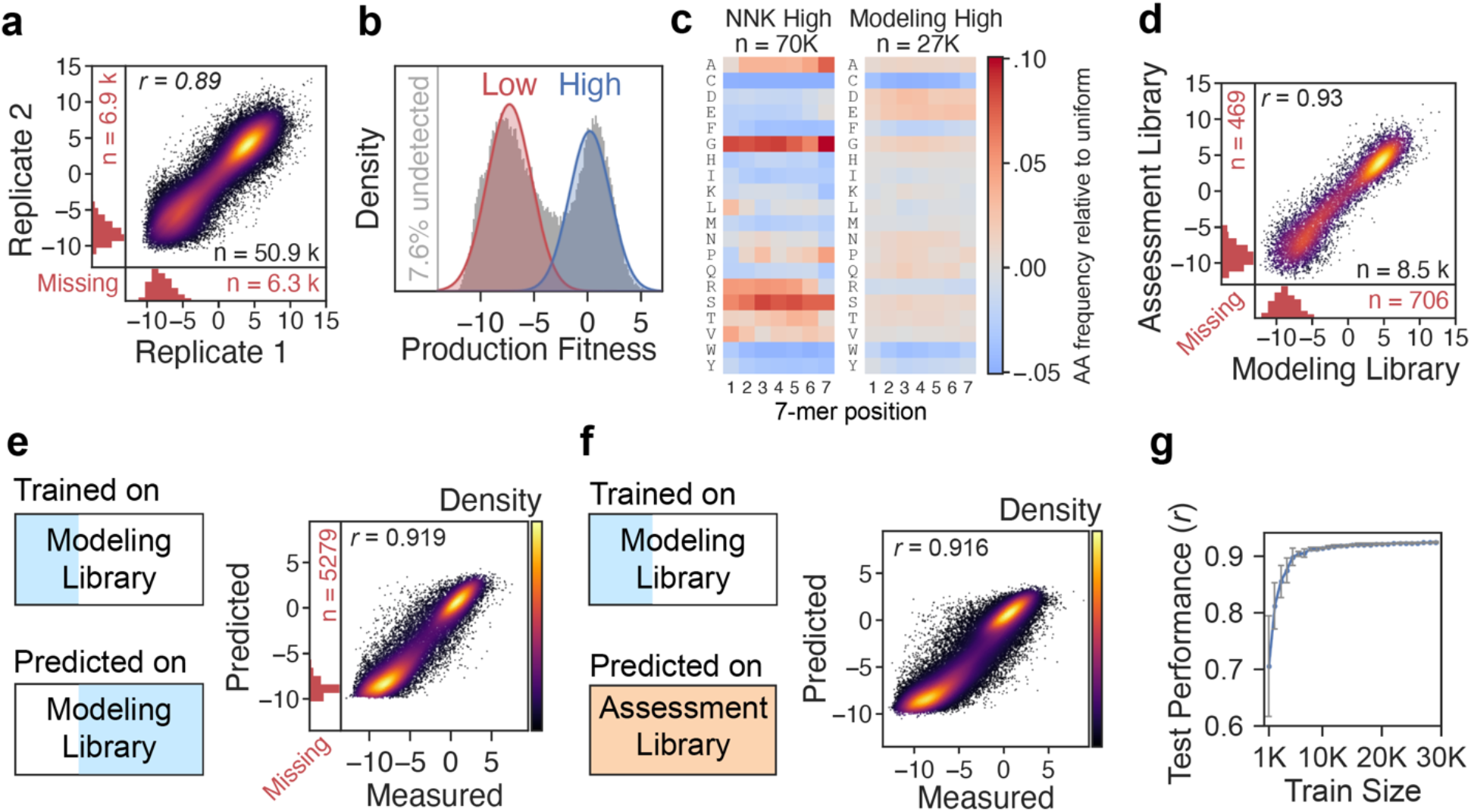
Mapping and learning the 7-mer production fitness landscape. (**a**) Correlation between the production fitness score of 7-mer replicate pairs. Each pair is aggregated across 12 replicates. The vertical and horizontal marginal histograms correspond to ‘missing’ cases, where only one 7-mer replicate of a pair was detected. (**b**) The production fitness distribution of the modeling library represents the variants detected in at least one of the 24 replicates (92.4% of total variants). The distributions representing low versus high production fitness are depicted. (**c**) The amino acid distribution by position for the variants in the 70K most abundant sequences in an NNK library versus the high production fitness distribution of the modeling library (27K out of 74.5K). (**d**) Production fitness replication quality of the control set (10K) that is shared between the modeling and assessment libraries. (**e, f**) Measured versus predicted production fitness score when the model is trained on a subset of the modeling library and tested on another subset of the same library (E) versus when tested on the independent assessment library, not including the overlapping 10K set (F). (**g**) Performance of the production fitness prediction model across different training set sizes (*n* = 10 models, mean ± s.d.).

The production fitness distribution of the modeling library could be modeled by a mixture of two Gaussian distributions: a “low fitness” versus a “high fitness” distribution (Fig. 2b). The low fitness distribution overlaps with the production fitness distribution of the stop codon-containing variants, which are presumably detected in the virus library due to cross-packaging (Supplementary Fig. 3). The variants in the high production fitness distribution exhibit distinguishing amino acid sequence characteristics, such as a general enrichment of negatively charged residues and depletion of cysteine and tryptophan (Fig. 2c). Nonetheless, this high production fitness distribution had less bias than an analogous set of the most abundant 70K variants from an NNK library (Fig. 2c). The production fitness scores for the 10K variants common to both libraries were consistent across the modeling and assessment libraries, suggesting that variant fitness is not noticeably impacted by the other variants in the libraries (Fig. 2d).

### A generalizable production fitness model

While prior studies applied classification models to predict AAV capsid production fitness ^14,15^, we used a regression model to capture the large variation in relative production fitness scores (±5-fold, log2 enrichment) within the high fitness and low fitness distributions. We first trained the model using the sequence and production fitness measurements of 24K variants unique to the modeling library. The accuracy of each model in this study was assessed by the agreement (Pearson correlation) between the measured fitness scores and the model’s predicted scores. Remarkably, the sequence-to-production-fitness model achieved high accuracy on the remaining subset of the library not used in the training process (Fig. 2e), as well as the independent assessment library (Fig. 2f). In addition, the model does not require large amounts of training data to obtain high accuracy, reducing the training from 24K to 5K variants only slightly reduced performance (*r* = 0.924±0.001 vs *r* = 0.899±0.015, Fig. 2g). These data demonstrate that the model is generalizable across libraries and to unseen variants and requires relatively small training datasets.

### Fit4Function enables reproducible data and accurate prediction models

Using the production fitness model, we randomly generated and predicted the fitness of 24M variants *in silico*. The predicted high production fitness sequence space was then evenly sampled for 240K variants to create a “Fit4Function” library (Fig. 3a). As expected, the measured fitness scores for the Fit4Function variants, when synthesized, mapped to a single distribution that closely follows the high production fitness distribution of the modeling library (Fig. 3a). The amino acid distribution in the Fit4Function library is similar to that of the high production fitness distribution from the modeling library and is similarly less biased when compared to that of the 240K most abundant variants in an NNK library (Fig. 3b). It is important to note that the criterion for high production fitness in populating Fit4Function libraries is not so stringent as to eliminate potentially promising functional candidates for downstream optimization; only variants with poor production (i.e., those whose production fitness is comparable to stop-codon containing control sequences) are considered low in production fitness and not sampled for Fit4Function libraries (Supplementary Fig. 3).

**Fig. 3.**
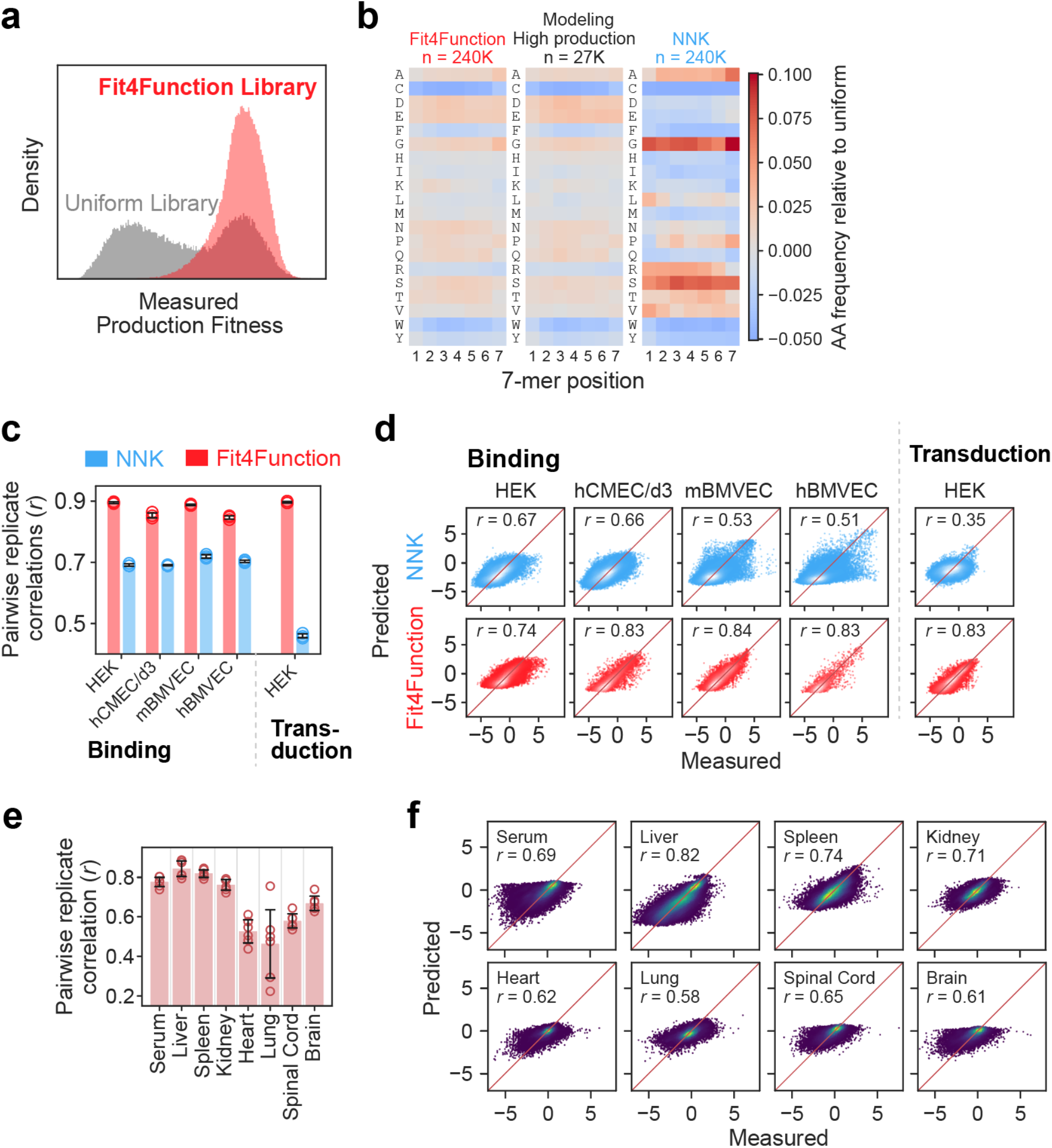
Fit4Function libraries evenly sample the high production fitness space and enable more accurate functional screening and prediction. (**a**) Distribution of the measured functional fitness scores for the Fit4Function library versus the uniform modeling library. (**b**) The amino acid distribution by position for the variants in the Fit4Function library, high production fitness distribution of the modeling library, and 240K most abundant sequences in an NNK library. (**c**) Quantitative comparison of pairwise Pearson correlations among biological triplicates for functional screens (mean ± s.d.; one-tailed paired t-test, *n* = 5 screens; *p* = 0.0074) using the Fit4Function library (240K) versus an NNK library (top 240K variants). hCMEC/d3: human brain endothelial cell line, mBMVEC: primary mouse brain microvascular endothelial cells, hBMVEC: primary human brain microvascular endothelial cells. (**d**) Measured versus predicted functional fitness (log2 enrichment scores) for models trained on Fit4Function versus NNK library data. (**e**) The replication quality (mean ± s.d.) between pairs of animals (*n* = 6 pairs across 4 animals) for the Fit4Function library biodistribution in eight organs. (**f**) The prediction performance of models trained on the *in vivo* biodistribution of the Fit4Function library across eight organs.

Fit4Function libraries are designed to enable the generation of reproducible and ML-compatible functional screening data. Specifically, the library is limited to a moderate size that enables deeper sequencing depth and samples only variants with high production fitness, which both enable more quantitative and reliable detection of each variant in the library. In addition, the library evenly samples the high production fitness amino acid sequence space, which results in less biased ML models that generalize well across the sequence space.

We compared the outcomes of our screening strategy using the Fit4Function library versus an NNK library across five functional assays: (1) HEK293 cell binding, (2) primary mouse brain microvascular endothelial cell (BMVEC) binding, (3) primary human BMVEC binding, (4) human brain endothelial cell line (hCMEC/D3) binding, and (5) HEK293 transduction. Binding and transduction were measured by quantitative sequencing capsid variant abundance at the DNA and mRNA levels, respectively. The Fit4Function library consistently yielded higher replication quality data than the NNK library (one-tailed paired t-test, *n* = 5 assays; *p* = 0.0074; Fig. 3c). We built and compared models trained on functional data derived from the Fit4Function library versus an NNK library (only data from the most abundant 240K variants in the NNK virus library was used). The Fit4Function-based models consistently achieved higher prediction accuracy (Fig. 3d).

We next sought to examine the use of the Fit4Function library to train prediction models of *in vivo* AAV biodistribution after systemic administration in adult C57BL/6J mice. The replication quality was high in liver, kidney, and spleen, and moderate in the brain, spinal cord, serum, heart, and lungs (Fig. 3e and Supplementary Fig. 4). We trained independent models to predict the variant tropism for each organ. The training data measurements were aggregated across three animals, and the data from the fourth animal was held out for independent testing. The models performed reasonably well when trained on assays with more reproducible data (Fig. 3f; model performance correlated with the data replication quality Fig. 3e), demonstrating the applicability of our approach to *in vivo* data.

**Fig. 4.**
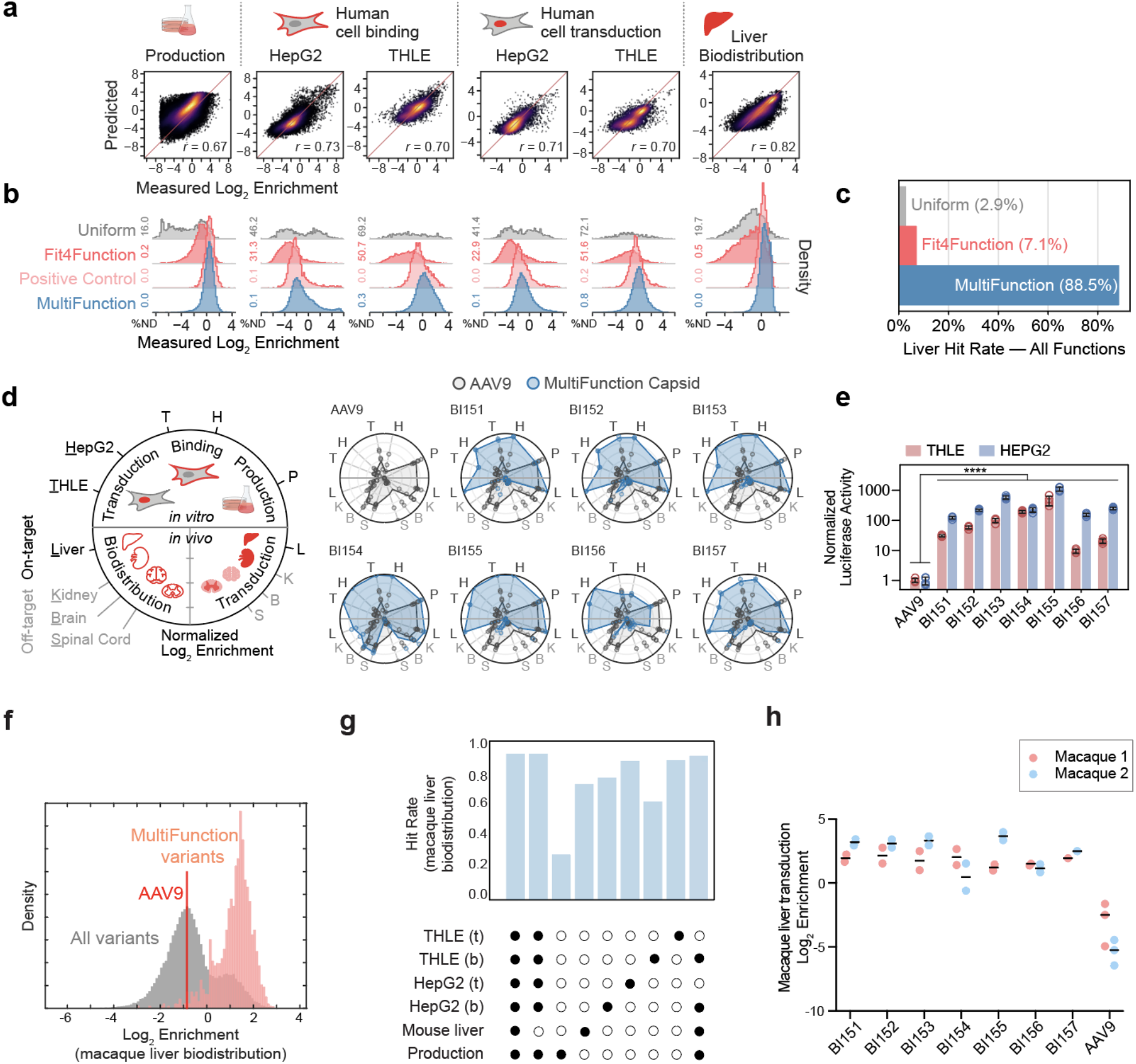
Liver-targeted MultiFunction library design, validation in human cells and mice, and translation to macaque. (**a**) The Pearson correlation of measured versus predicted enrichment for production fitness and functional assays relevant to hepatocyte cross-species targeting. (**b**) The distribution of enrichment across variants sampled from the Uniform (3K), Fit4Function (10K), Positive Control (Fit4Function variants satisfying the six conditions), and MultiFunction libraries. The histograms are density-normalized, including non-detected variants (ND). (**c**) The hit rate for variants satisfying the six conditions in each listed variant set. Positive Control variants were sampled from the Fit4Function variants and by design meet all six conditions; hence, these are not plotted. (**d**) On-target and off-target measurements for the seven selected capsids (BI151–157) and AAV9 in the MultiFunction library pool, shown as normalized log2 enrichments of the selected capsid (two 7-mer replicates) as compared to AAV9 (four 7-mer replicates). Measured enrichment was linearly normalized according to the maximum and minimum enrichment values for each assay across all capsids. Individual 7-mer replicates are plotted as points, and the average normalized enrichments across replicates are plotted as polygon vertices. (**e**) HepG2 and THLE transduction were assessed 24 hours post-transduction at 3000 vg/cell using a luciferase assay (*n* = 4 transduction replicates per group, mean ± s.d., \*\*\*\**p*<1e-4, unpaired, one-sided t-tests on log-transformed values, and Bonferroni corrected for multiple-hypotheses). Luciferase relative light units were normalized to AAV9. (**f**) A 100K variant Fit4Function library was injected intravenously into a cynomolgus macaque and biodistribution was assessed four hours later. Variants in the administered library predicted to concurrently meet the six trait conditions were observed to be highly enriched in terms of macaque liver biodistribution. The density plot shows the distribution of variants normalized to the sum of counts for each indicated set of variants. (**g**) The fraction of the indicated MultiFunction variants that are enriched in the macaque liver (defined as at least two-fold log2 enrichment greater than that of AAV9) are shown for each combination of predicted traits. Binding and transduction are indicated by ‘b’ and ‘t’, respectively. For instance, of the MultiFunction variants that were predicted to exhibit all six traits, 91.1% were found to be enriched in the cynomolgus macaque liver. (**h**) The macaque liver transduction efficiency for the seven individually characterized liver MultiFunction variants are shown (*n* = 2 rhesus macaques). In the virus library, each variant was represented by two 7-mer replicates while AAV9 was represented by three replicates.

### Multi-trait capsid identification

Efficient and durable gene delivery to the liver remains challenging due to vector efficiency and capsid antigen presentation and T cell-mediated immunity. Liver-directed therapies should benefit from the development of more potent AAV vectors that can be administered at lower doses to reduce the exposure to capsid antigens. Previous efforts to develop capsids with improved human hepatocyte transduction generated candidates that are selective for human hepatocytes but inefficiently transduced the mouse liver *in vivo*^2,16^. While such vectors have important translational potential – one capsid, LK03 is now being evaluated in human clinical trials (NCT05092685; NCT04581785; NCT03003533; NCT03876301)^2^ – there is a need for capsids that are also compatible with preclinical efficacy and safety testing. Our objective was to design a ‘MultiFunction’ library consisting only of variants that are each predicted to possess multiple enhanced functions related to crossspecies hepatocyte gene delivery. Toward this goal, we performed five separate functional screens of the Fit4Function library: (1) binding or (2) transduction of the human hepatocellular carcinoma cell line (HepG2), (3) binding or (4) transduction of the human liver epithelial cell line (THLE), and (5) efficient liver biodistribution in C57BL/6J mice (Supplementary Fig. 5). We used the high-quality data from these functional screens to train and assess the performance of five independent sequence-to-function models (Fig. 4a). With the production fitness model and these five functional fitness models, we screened 10M randomly generated capsid variants *in silico* and selected 30K liver-targeted MultiFunction candidate variants predicted to have enhanced phenotypes across all five functions and production fitness (“enhanced phenotype” was arbitrarily defined as any variant above the 50th percentile of the measured enrichment scores). In the MultiFunction library, each variant was encoded by two nucleotide sequences serving as biological replicates. In addition, we included 3K variants from the modeling library (high and low production fitness; Uniform Control), 10K from the Fit4Function library (Fit4Function Control), and 3K from the known hits in the Fit4Function library, i.e., variants from the Fit4Function library that had been experimentally confirmed to exhibit enhanced phenotypes for the five hepatocyte-related traits and production fitness (Positive Control).

To assess the accuracy of our predictions and identify the top-performing variants, we screened the MultiFunction library on the same five assays related to hepatocyte targeting and on production fitness (Supplementary Fig. 6). The MultiFunction variants either matched or surpassed the performance of the positive controls from the Fit4Function library (Fig. 4b); 88.5% of the MultiFunction library variants satisfied our enhanced phenotype definition as compared to 2.9% of sequences in the uniform space or 7.1% of the Fit4Function library control (Fig. 4c). Although the 7-mer sequences in the MultiFunction library have an increased frequency of arginines and lysines, the library diversity remains high (Supplementary Fig. 6d).

We individually assessed the performance of seven variants that were selected from the MultiFunction library based on their measured production fitness, liver biodistribution and transduction in mice, and their enhanced ability to bind and transduce human HEPG2 and THLE cells (Fig. 4d). Each capsid and AAV9, as a control, were used to package a single-stranded green fluorescent protein (GFP) and Luciferase dual reporter AAV2 genome. Production yields were comparable to that of AAV9 (Supplementary Fig. 7a). When administered to mice at 1×10^10^ vg/mouse and assessed for GFP expression three weeks later, each capsid and AAV9 efficiently transduced hepatocytes as assessed by the native GFP fluorescence in DAPI^+^ liver nuclei (Supplementary Fig. 7b, c and 8). All novel AAVs were more effective (10 to 1000-fold) than AAV9 at transducing the HEPG2 and THLE cell lines (Fig. 4e and Supplementary Fig. 7d).

### Fit4Function translates across species to macaques

We administered a 100K member Fit4Function library intravenously to an adult cynomolgus macaque and assessed biodistribution. Liver-targeted MultiFunction capsids, predicted with the six prior models that were trained only on human cell and mouse data and production fitness, were highly enriched in terms of macaque liver biodistribution (Fig. 4f). The combination of multiple functional predictors was more effective at identifying variants with increased biodistribution to the macaque liver than any single predictor used in isolation (Fig. 4g). The five liver models exhibited redundancy, which is unsurprising given that they are readouts of related functions (Fig. 4g). Notably, the *in vitro* human hepatocyte transduction models translated better to cynomolgus macaque liver biodistribution compared to the *in vivo* mouse liver biodistribution model, which was neither necessary nor sufficient to demonstrate transferability to cynomolgus macaque liver biodistribution; the hit rate did not decrease when the mouse liver model was excluded from the combination of models (Fig. 4g). The hit rate decreased only modestly when both human hepatocyte transduction models were excluded, demonstrating the utility of using models in combination (Fig. 4g). All seven of the liver MultiFunction capsids individually validated in human cells and in mice *in vivo* (Fig. 4d, Supplementary Fig. 7) were more efficient than AAV9 at transducing the rhesus macaque liver when administered as a library (Fig. 4h; *n* = 2 rhesus macaques).

## Discussion

The Fit4Function pipeline presents a significant conceptual and technological advance over prior AAV engineering studies, including those that leverage ML. Conventional *in vivo* selections use sequential rounds to narrow the focus of sequence exploration to a handful of top candidates, which may not have other traits required for translation to preclinical models and clinical trials. Simultaneously engineering multiple traits into AAV capsids or other proteins of interest is an important but challenging goal. To date, most protein engineering efforts, including those leveraging ML, have focused on optimizing a single function, e.g. generating more efficiently produced and diversified AAV capsid libraries but stopping short of multi-trait prediction ^14,18,19^. A few groups have gone beyond single trait engineering by combining multiple previously validated functional structures into a single protein, e.g., by recombining structurally independent segments from different channelrhodopsins possessing known functions, localizations, and photocurrent properties of interest ^20^, or by applying protein design tools to filter out variants that do not meet additional characteristics such as solubility and immunogenicity ^21^. However, as these strategies rely on the recombination of multiple existing functional structures into a single protein or the use of third-party protein design tools, they cannot be broadly generalized to engineer multiple *de novo* functions. A key obstacle to combining multiple ML models that predict different traits is the aggregated error that increases with each added model. The Fit4Function approach directly tackles this problem by leveraging a moderately sized, all viable, low-bias (ML-designed) library to generate highly reproducible data for multi-trait learning with a low false positive rate. This allows the models to be applied in different combinations with a low risk of aggregating significant error. MultiFunction libraries can thus be generated to more efficiently explore the vast sequence space for multi-trait capsids.

The Fit4Function approach can help to reduce the need for extensive screening in animals in two ways. Firstly, the unique features of Fit4Function libraries enable the quantitative assessment of capsid biodistribution and candidate selection from just a single round of screening. It is only necessary to screen a Fit4Function library once for a given function to then predict the functionality of sequences that were not contained in the original library. In contrast, it typically requires two or more rounds of *in vivo* screening to reliably identify top candidates from conventional selections, and the data from these screens cannot be used to accurately predict the traits of variants not tested in that screen. This means that the Fit4Function approach can be used to design libraries full of diverse and promising candidates for more efficient screening in animals or *in vitro* assays. Secondly, unlike existing screening strategies, our approach can systematically determine the functional assays or combinations thereof that drive cross-species transferability. As the Fit4Function approach is applied to more functions of interest (e.g., crossing the blood-brain barrier), it will become apparent whether it is worthwhile to continue screening in mice or other animals for those functions. This can inform the choice of cell or animal models to perform screens in and develop vectors that are more likely to translate preclinically and clinically.

As with other ML-guided approaches, Fit4Function can be more challenging to implement with assays that produce low quality data due to lower detection sensitivities. For example, data reproducibility and subsequent model performance can be bottlenecked by *in vivo* transduction assays in some organs due to the inherent tropism of the parental capsid, inter-animal variability, and technical challenges related to tissue sampling. One approach to improve data quality with low sensitivity assays may be to use smaller Fit4Function libraries because reducing library diversity increases the sampling of each individual variant and therefore the quality of the screening data. A second limitation that affects any multi-objective engineering effort is that variants that are maximally optimized for multiple objectives may not exist, especially in cases where performance on functions are negatively correlated. While Fit4Function cannot overcome this fundamental problem, it provides the means to efficiently search the vast production fit sequence space for variants that are reasonably well optimized for multiple traits.

With continued application across experiments and laboratories, the Fit4Function approach should enable the assembly of a vast ML atlas that can accurately predict the performance of AAV capsid variants across dozens of traits and inform the design of screening pipelines. In addition, the Fit4Function approach should translate to engineering other proteins that are amenable to quantitative, high-throughput screening of libraries that are diversified at a defined set of residues.

## Methods

### Modeling and assessment library design

The modeling and assessment libraries were designed to contain 150K nucleotide sequences each. The libraries were composed of 64.5K unique and 10K shared amino acid sequences generated by uniformly sampling all 20 amino acids at each position. The 74.5K variants were duplicated via 7-mer replication. 1K sequences containing stop codons were included to detect problems with cross packaging. In total, each library comprised a final set of 150K sequences.

### Capsid library synthesis

To produce synthetic library inserts, lyophilized DNA oligonucleotide libraries (Agilent G7223A) or NNK hand mixed primers (IDT) were spun down at 8000 RCF for 1 minute, resuspended in 10 μL UltraPure DNase/RNase-Free Distilled Water (ThermoFisher Scientific, 10977015), and incubated at 37°C for 20 minutes. For pooled synthetic oligonucleotide libraries, the following primer format was used: 5’-GTATTCCTTGGTTTTGAACCCAACCGGTCTGCGCCTGTGC-(NNN)7- TTGGGCACTCTGGTGGTTTGTGGCCAC. To produce NNK inserts, the AAV9_K449R_Forward (5’-CGGACTCAGACTATCAGCTCCC) and AAV9_K449R_NNK_Reverse (GTATTCCTTGGTTTTGAACCCAACCGGTCTGCGCCTGTGC-(MNN)7- TTGGGCACTCTGGTGGTTTGTG) primers were used.

To amplify the oligonucleotide libraries and incorporate them into an AAV9 (K449R) template, 2 μL of the resuspended pooled oligonucleotide library or NNK-based library was used as an initial reverse primer along with 0.5 μM AAV9_K449R_Forward primer in a 25 μL PCR amplification reaction using Q5 Hot Start High-Fidelity 2X Master Mix (NEB, M0494S). 50 ng of a plasmid containing only AAV9 (K449R) VP1 amino acids 347-586 was used as a PCR template. PCR was performed following the manufacturer’s protocol with an annealing temperature of 65°C for 20 seconds and an extension time of 90 seconds. After six PCR cycles, 0.5 μM AAV9_K449R_Reverse (5’-GTATTCCTTGGTTTTGAACCCAACCG) was spiked into the reaction as a reverse primer to further amplify sequences containing the oligonucleotide library for an additional 25 cycles. To remove the PCR template, 1 μL of DpnI (NEB, R0176S) was added to the PCR reaction and incubated at 37°C for one hour. Afterwards, the PCR products were cleaned using AMPure XP beads (Beckman, A63881) following the manufacturer’s protocol.

The PCR insert was assembled into 1600 ng of a linearized mRNA selection vector (AAV9-CMV-Express) with NEBuilder HiFi DNA Assembly Master Mix (NEB, E2621L) at a 3:1 insert:vector Molar ratio in a 80 μL reaction volume, incubated at 50°C for one hour, and then at 72°C for 5 minutes. Afterwards, 4 μL of Quick CIP (NEB, M0508S) was spiked into the reaction and incubated at 37°C for 30 minutes to dephosphorylate unincorporated dNTPs that may inhibit downstream processes. Finally, 4 μL of T5 Exonuclease (NEB M0663S) was added to the reaction and incubated at 37°C for 30 minutes to remove unassembled products. The final assembled products were cleaned using AMPure XP beads (Beckman, A63881) following the manufacturer’s protocol and their concentrations were quantified with a Qubit dsDNA HS Assay Kit (ThermoFisher Scientific, Q32851) and a Qubit fluorometer.

### mRNA selection vector

The mRNA selection vector (AAV9-CMV-Express) was designed to enrich for functional AAV capsid sequences by recovering capsid mRNA from transduced cells. AAV9-CMV-Express uses a ubiquitous CMV enhancer and AAV5 p41 gene regulatory elements to drive AAV Cap expression. The AAV-Express plasmid was constructed by cloning the following elements into an AAV genome plasmid in the following order: a cytomegalovirus (CMV) enhancer-promoter, a synthetic intron and the AAV5 P41 promoter along with the 3’ end of the AAV2 Rep gene, which includes the splice donor sequences for the capsid RNA. The capsid gene splice donor sequence in AAV2 Rep was modified from a non-consensus donor sequence CAGGTACCA to a consensus donor sequence CAGGTAAGT. The AAV9 capsid gene sequence was synthesized with nucleotide changes at S448 (TCA to TCT, silent mutation), K449R (AAG to AGA), and G594 (GGC to GGT, silent mutation) to introduce restriction enzyme recognition sites for oligonucleotide library fragment cloning. The AAV2 polyadenylation sequence was replaced with a simian virus 40 (SV40) late polyadenylation signal to terminate the capsid RNA transcript.

### Virus production

For library production, HEK293T/17 cells (ATCC, CRL-11268) were seeded at 22 million cells per 15 cm plate the day before transfection and grown in DMEM with GlutaMAX (Gibco, 10569010) supplemented with 5% FBS and 1X non-essential amino acid solution (NEAA) (Gibco, 11140050). The next day, each plate was triple transfected with 39.93 μg of total plasmid DNA encoding pHelper, RepStop encoding the AAV2 Rep genes, pUC19 at a ratio of 2:1:1, respectively, and with 10 ng of assembled library DNA. The media was exchanged for fresh DMEM with 5% FBS and 1X NEAA at 20 hours post transfection. At 60 hours, the media and cell lysates were harvested and purified following the published protocol ^22^.

Individual recombinant AAVs were produced in suspension HEK293T cells, using F17 media (Thermofisher). Cell suspensions were incubated at 37°C, 8% CO2, 125 RPM. 24 hours before transfection, cells were seeded in 200 mL at ~1 million cells/mL. The day after, cells (~2 million cells/mL) were transfected with pHelper, pRepCap and pTransgene (2:1:1 ratio, 2 μg DNA per million cells) using Transport 5 transfection reagent (Polysciences) with a 2:1 PEI:DNA ratio. Three days post-transfection, cells were pelleted at 2000 RPM for 10 minutes into Nalgene conical bottles. The supernatant was discarded, and cell pellets were stored at −20°C until purification. Each pellet, corresponding to 200 mL of cell culture, was resuspended in 7 mL of 500 mM NaCl, 40 mM Tris-base, 10 mM MgCl2, with Salt Active Nuclease (ArcticZymes, #70920-202) at 100 U/mL. Afterwards, the lysate was clarified at 2000 RCF for 10 minutes and loaded onto a density step gradient containing OptiPrep (Cosmo Bio, AXS-1114542) at 60%, 40%, 25%, and 15% at a volume of 5, 5, 6, and 6 mL respectively in OptiSeal tubes (Beckman, 361625). The step gradients were spun in a Beckman Type 70ti rotor (Beckman, 337922) in a Sorvall WX+ ultracentrifuge (Thermo Scientific, 75000090) at 69,000 RPM for 1 hour at 18°C. Afterwards, ~4.5 mL of the 40-60% interface was extracted using a 16-gauge needle, filtered through a 0.22 μm PES filter, buffer exchanged with 100K MWCO protein concentrators (Thermo Scientific, 88532) into PBS containing 0.001% Pluronic F-68, and concentrated down to a volume of 500 μL. The concentrated virus was filtered through a 0.22 μm PES filter and stored at 4°C or −80°C.

### AAV Titering

To determine AAV titers, 5 μL of each purified virus library were incubated with 100 μL of an endonuclease cocktail consisting of 1000U/mL Turbonuclease (Sigma T4330-50KU) with 1X DNase I reaction buffer (NEB B0303S) in UltraPure DNase/RNase-Free distilled water at 37°C for one hour. Next, the endonuclease solution was inactivated by adding 5 μL of 0.5M EDTA, pH 8.0 (ThermoFisher Scientific, 15575020) and incubated at room temperature for 5 minutes and then at 70°C for 10 minutes. To release the encapsidated AAV genomes, 120 μL of a Proteinase K cocktail consisting of 1M NaCl, 1% N-lauroylsarcosine, 100 μg/mL Proteinase K (Qiagen, 19131) in UltraPure DNase/RNase-Free distilled water was added to the mixture and incubated at 56°C for 2 to 16 hours. The Proteinase K-treated samples were then heat-inactivated at 95°C for 10 minutes. The released AAV genomes were serial diluted between 460-460,000X in dilution buffer consisting of 1X PCR Buffer (Thermo Fisher Scientific, N8080129), 2 μg/mL sheared salmon sperm DNA (Thermo Fisher Scientific, AM9680), and 0.05% Pluronic F68 (Thermo Fisher Scientific, 24040032) in UltraPure Water (Thermo Fisher Scientific). 2 μL of the diluted samples were used as input in a ddPCR supermix (Bio-Rad, 1863023). Primers and probes, targeting the ITR and CAG promoter region, were used for titration, at a final concentration of 900 nM and 250 nM, respectively (ITR2_Forward: 5’-GGAACCCCTAGTGATGGAGTT; ITR2_Reverse: 5’-CGGCCTCAGTGAGCGA; ITR2_Probe: 5’-CACTCCCTCTCTGCGCGCTCG [FAM/Iowa Black FQ Zen]; CAG_Forward: 5’-TGTTCCCATAGTAACGCCAATAG; CAG_Reverse: GTACTTGGCATATGATACACTTGATG; CAG_Probe: 5’-TTACGGTAAACTGCCCACTTGGCA [FAM/Iowa Black FQ Zen]). Droplets were generated using a QX100 Droplet Generator following the manufacturer’s protocol. The droplets were transferred to thermocycler and cycled according to the manufacturer’s protocol with an annealing/extension of 58°C for one minute. Finally, droplets were read on a QX100 Droplet Digital System to determine titers.

### Assessing production fitness

To recover only encapsidated AAV genomes for downstream analysis, 10^11^ viral genomes were extracted using the endonuclease and Proteinase K steps outlined above (AAV Titering). After Proteinase K treatment, samples were column purified using a DNA Clean and Concentrator Kit (Zymo Research, D4033) and eluted in 25 μL elution buffer for NGS preparation.

### NGS sample preparation

To prepare AAV libraries for sequencing, qPCR was performed on extracted AAV genomes or cDNA to determine the cycle thresholds for each sample type to prevent overamplification. PCR amplification using equal primer pairs (1-8) (described in Huang et al. 2022) ^23^ was used to attach partial Illumina Read 1 and Read 2 sequences using Q5 Hot Start High-Fidelity 2X Master Mix with an annealing temperature of 65°C for 20 seconds and an extension time of 60 seconds. Round one PCR products were purified using AMPure XP beads following the manufacturer’s protocol and eluted in 25 μL UltraPure Water (Thermo Fisher Scientific). 2 μL was used as input in a second round of PCR to attach on Illumina adaptors and dual index primers (NEB, E7600S) for five PCR cycles using Q5 HotStart-High-Fidelity 2X Master Mix with an annealing temperature of 65°C for 20 seconds and an extension time of 60 seconds. The round two PCR products were purified using AMPure XP beads following the manufacturer’s protocol and eluted in 25 μL UltraPure DNase/RNase-Free distilled water (Thermo Fisher Scientific).

To quantify the amount of PCR products for NGS, an Agilent High Sensitivity DNA Kit (Agilent, 5067-4626) was used with an Agilent 2100 Bioanalyzer. PCR products were pooled and diluted to 2-4 nM in 10 mM Tris-HCl, pH 8.5 and sequenced on an Illumina NextSeq 550 following the manufacturer’s instructions using a NextSeq 500/550 Mid or High Output Kit (Illumina, 20024904 or 20024907), or on an Illumina NextSeq 1000 following the manufacturer’s instructions using NextSeq P2 v3 kits (Illumina, 20046812). Reads were allocated as follows: I1: 8, I2: 8, R1: 150, R2: 0.

### NGS data processing

Sequencing data was de-multiplexed with *bcl2fastq* (version v2.20.0.422) using the default parameters. The Read 1 sequence (excluding Illumina barcodes) was aligned to a short reference sequence of AAV9: CCAACGAAGAAGAAATTAAAACTACTAACCCGGTAGCAACGGAGTCCTATGGACAAGTGGCCACAAACCACCAGAGTGCCCAANNNNNNNNNNNNNNNNNNNNNGCACAGGCGCAGACCGGTTGGGTTCAAAACCAAGGAATACTTCCG Alignment was performed with *bowtie2* (version 2.4.1) ^24^ with the following parameters: --end-to-end --very-sensitive --np 0 --n-ceil L,21,0.5 --xeq-N 1 --reorder --score-min L,-0.6,-0.6 −5 8 −3 8

Resulting sam files from *bowtie2* were sorted by read and compressed to bam files with *samtools* (version 1.11- 2-g26d7c73, *htslib* version 1.11-9-g2264113) ^*25,26*^.

*Python* (version 3.8.3) scripts and *pysam* (version 0.15.4) were used to extract the 21 nucleotide insertion from each amplicon read. Each read was assigned to one of the following bins: Failed, Invalid, or Valid. Failed reads were defined as reads that did not align to the reference sequence, or that had an in/del in the insertion region (i.e., 20 bases instead of 21 bases). Invalid reads were defined as reads whose 21 bases were successfully extracted, but matched any of the following conditions: 1) Any one base of the 21 bases had a quality score (AKA Phred score, QScore) below 20, i.e., error probability > 1/100, 2) Any one base was undetermined, i.e., “N”, 3) The 21 base sequence was not from the synthetic library (this case does not apply to NNK library). Valid reads were defined as reads that did not fit into either the Failed or Invalid bins. The Failed and Invalid reads were collected and analyzed for quality control purposes, and all subsequent analyses were performed on the Valid reads.

Count data for valid reads was aggregated per sequence, per sample, and was stored in a pivot table format, with nucleotide sequences on the rows, and samples (Illumina barcodes) on the columns. Sequences not detected in samples were assigned a count of 0.

### Data normalization

Count data was read-per-million (RPM) normalized to the sequencing depth of each sample (Illumina barcode) with:

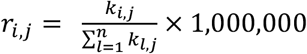

Where *r* is the RPM-normalized count, *k* is the raw count, *i* = *1… n* sequences, and *j* = *1… m* samples.

As each biological sample was run in triplicate, we aggregated data for each sample by taking the mean of the RPMs

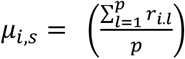

across p replicates of sample s. We estimated normalized variance across replicates by taking the coefficient of variation (*CV*):

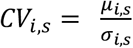

where *sigma{i,s}* is the standard deviation for variant *i* in sample *s* over *p* replicates.

Log_2_ enrichment for each sequence was defined as:

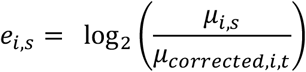

Where *e* is the log_2_ enrichment, *mu* is the mean of the replicate RPMs, and *t* is the normalization sample. For production fitness, the sample *s* is the variant abundance after virus production, and the normalization sample *t* is the variant abundance in the plasmid pool. For functional screens, the sample *s* is the variant abundance of the screen, and the normalization factor *t* is the variant abundance after virus production. To avoid dividing by 0 in *e* (for NNK library processing), *mu_corrected* is defined as:

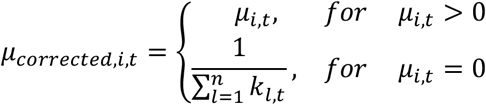

I.e., counts of 0 across all 3 replicates for the normalization sample were adjusted to a count of 1 across all 3 replicates.

### Production fitness training and assessment

We designed a robust ML framework that we used for the production fitness and Fit4Function functional mappings. A long short-term memory (LSTM) regression model with two hidden layers of 140 and 20 nodes was implemented in *Keras ^27^*. RNNs, and LSTMs in particular, have been successfully applied for learning functions from biological sequence data as they are designed to capture local and distant relationships across different parts of the input sequences ^14,28^. Model parameters and hyperparameters were subject to fine tuning processes but no significant performance was gained across all different functional models implemented in this study. Therefore, we kept a simple model architecture across all modeling throughout this study. The input layer was 7-mer amino acid sequences one-hot encoded into a 20 × 7 matrix. The target/output is the relative production (or functional) fitness score. Loss was optimized by mean-squared-error with Adam optimizer running on a learning rate of 0.001 ^29^. The batch size was set to 500 observations. To avoid overfitting, model training was controlled by a custom early stopping procedure where the training process was terminated if the ratio of training error to validation error dropped below 0.90.

For production fitness learning, the training size was optimized by training the framework on increments of 1K variants. Variants that were not detected (*n* = 5,279) after virus production were filtered out from training. Model validation performance was reported at each training size, and a size of 24K variants was arbitrarily selected for final model training given that the model performance reached a plateau after a training size of ~5K. The modeling library core variants (N ~ 60K, after removing the non-detected sequences) were then randomly divided into training (24K), validation (12K) and testing subsets (24K), all from the modeling library. The model was trained on the training set (24K), validated during the training process on the validation set (12K), and tested on the testing set (24K). The model was further tested on the unique variants from the assessment library to assess its generalization across libraries.

### Fit4Function library sampling

The Fit4Function libraries are intended to be sampled from the high production fitness space. For the Fit4Function library utilized in this study, we first uniformly sampled a set of 7-mer amino acid sequences 100 times the required library size (240K Fit4Function variants * 100 = 24M variants), by equally sampling each amino acid at each of the 7 positions. Duplicates were removed and the remaining sequences were scored using the production fitness model. Then, the 240K Fit4Function library variants were probabilistically sampled from the parametrized high production fitness distribution. In addition to the 240K high production fitness variants, we added 1K stop codon-containing variants and 3K variants from the 10K shared variants between the modeling and assessment libraries as a control set.

### Fit4Function library validation

Fitness enrichment scores are relative across library variants due to normalization calculations; calibration is needed to make the fitness scores of two libraries of different compositions comparable for assessment or integration purposes. To calibrate the Fit4Function library production fitness, we used the 3K control set to fit an ordinary linear regression model of the measured production fitness scores between the Fit4Function library and the modeling library. These regression parameters were applied to the production fitness measured scores of the 240K Fit4Function variants to obtain calibrated production fitness scores. After synthesizing the Fit4Function library, we compared, by means of correlation, the predicted fitness scores to the calibrated measured fitness.

### Animals

All mouse procedures were performed as approved by the Broad Institute Institutional Animal Care and Use Committee (IACUC), approval number 0213-06-18-1. Female C57BL/6J (000664) mice were obtained from the Jackson Laboratory (JAX). Recombinant AAV vectors were administered intravenously via the retro-orbital sinus in young adult (7- to 8-week-old) animals (*n* = 5 mice per vector group). Mice were randomly assigned to groups based on predetermined sample sizes. No mice were excluded from the analyses. For all assays, mice were anesthetized with EUTHASOL™ (Virbac) and transcardially perfused with phosphate buffer saline, pH 7.4, at room temperature (RT). Experimenters were not blinded to the sample groups.

For the cynomolgus macaque experiment (*n* = 1), the study plan involving the care and use of animals was reviewed and approved by the Charles River CR-LAV Institutional Animal Care and Use Committee (IACUC). During the study, the care and use of animals was conducted by CR-LAV with guidance from the USA National Research Council and the Canadian Council on Animal Care (CCAC). The Test Facility is accredited by the CCAC and AAALAC. Per the CCAC guidelines, this study was considered as a category of invasiveness C.

The rhesus macaque study (*n* = 2) was conducted in the NIH Nonhuman Primate Testing Center for Evaluation of Somatic Cell Genome Editing Tools at the University of California, Davis. All procedures conformed to the requirements of the Animal Welfare Act, and protocols were approved prior to implementation by the UC Davis IACUC.

### AAV mouse *in vivo* biodistribution assays

Purified virus libraries were injected IV (retro-orbital sinus) at a dose of 1×10^12^ into C57BL/6J mice. Two hours post-injection serum was collected and organs were harvested using disposable 3 mm biopsy punches (Integra, 33-32-P/25) with a new biopsy punch used per organ per replicate. Harvested tissues were immediately frozen on dry ice. AAV genomes were recovered using a DNeasy kit (Qiagen, 69504) following the manufacturer’s protocol and samples were eluted in 200 μL elution buffer for NGS preparation.

### AAV cynomolgus macaque *in vivo* biodistribution assays

The library administered had 100K unique amino acid variants following the Fit4Function criteria (uniformly sampled from the high production fitness sequence space) in addition to a calibration set (3K), control variants, and AAV9. Each variant in the Fit4Function distribution was represented by either two or six 7-mer replicates; AAV9 was represented by two replicates. The purified virus library was injected IV at a dose of 4.6 x 10^12^ vg/kg into a 3.4 kg adult female cynomolgus macaque that was pre-screened for NAbs against AAV9 (CRL). Four hours after systemic delivery, the animal was perfused with cold PBS and organs were harvested and snap frozen on dry ice. DNA was extracted using a DNeasy kit in a Qiagen QIAcube Connect. Samples were then processed as detailed in the NGS sample preparation section.

### AAV rhesus macaque *in vivo* transduction assays

Approximately 3-month-old rhesus monkeys (~1 kg; one male, one female) were screened then assigned to the project after confirming seronegative status for AAV9 antibodies using standardized Testing Center assays. Sedation with Telazol (IM) was performed prior to IV administration of a purified virus library (1 x 10^13^ vg/kg) with blood samples collected (~4 mL; hematology, clinical chemistry, serum, plasma; pre-administration then weekly post-administration). Animals were monitored closely during the study period and until endpoint (four weeks post-administration). They remained robust and healthy with no evidence of adverse findings (e.g., daily physical signs, body weights, hematology and clinical chemistry panels were within the normative range at all timepoints; data not shown). Four weeks after systemic delivery, tissues were collected and snap frozen over liquid nitrogen then placed on dry ice immediately prior to storage at ≤-80°C. RNA and DNA were extracted using TRIzol (Invitrogen, 15596026) following the manufacturer’s instructions. Total RNA was then processed through a RNeasy kit (Qiagen, 74106) followed by on-column DNA digestion. RNA was converted to cDNA using Maxima H Minus Reverse Transcriptase (Thermo Scientific, EP0751) according to the manufacturer’s instructions. Samples were then processed as detailed in the NGS sample preparation section.

### Rhesus macaque serum screening for anti-AAV9 neutralizing antibodies

Neutralization assays were also performed at two MOIs, 500 and 1000, in Perkin-Elmer white 96-well plates. Four-fold serial dilutions (1:4 to 1:16,384) of macaque serum samples were prepared in 96-well plates using DMEM supplemented with 5% FCS. Then, 40 μL of each dilution was transferred to a separate 96-well plate, mixed with an equal volume of AAV9.CAG-GFP-P2A-Luciferase-WPRE-SV40 vector (4–8E7 vg per 40 μL, diluted in DMEM-5% fetal bovine serum), and incubated for one hour at 37°C. Following the incubation, AAV-serum samples were transferred into a new 96-well plate (20 μL triplicates) and a total of 80 μL of DMEM-5% fetal bovine serum, containing 20,000 HEK293T cells, was added to each well (final volume of 100 μL). 96- well plates were incubated for 48 hours at 37°C, 5% CO2. Luminescence levels were read using a Perkin Elmer Victor Luminescence Plate Reader using the britelite plus Reporter Gene Assay System (Perkin-Elmer, #6066761). Data was analyzed using the *neutcurve* Python package developed by the Bloom laboratory. The neutralizing antibody titer was measured as the concentration that resulted in a 50% reduction in luciferase activity relative to the no-serum control. Animals used in the transduction study had NAb titers <1:12 in this set of antibody screens.

### *In vitro* binding and transduction

HEK293T/17 (ATCC^®^ CRL-11268™), HepG2 (ATCC^®^ HB-8065™), THLE-2 (ATCC^®^ CRL-2706™), hCMEC/D3 (Millipore, SCC066), and human and mouse BMVECs (Cell Biologics, H-6023 and C57-H6023) were grown in 100 mm dishes and exposed to the Fit4Function or (NNK) 7-mer library (MOI of 1E4 for HEK293T/17, MOI of 3E4 for hCMEC/D3, MOI of 6E4 for primary human and mouse BMVECs, and MOI of 5E3 for HepG2 and THLE-2) diluted in 10 mL of growth media at 4°C with gentle rocking for two hours. Cells were then washed three times with DPBS, and total DNA was extracted with the DNeasy kit (Qiagen) according to the manufacturer’s instructions. Half of the recovered DNA was used in PCR amplification for viral genome sequence recovery.

Transduction assays were performed as described above with the following exceptions: The cells were cultured in growth media containing virus for 60 hours and total RNA was then extracted with the RNeasy kit (Qiagen). From the total RNA, 5 μg was converted to cDNA using the Maxima H Minus Reverse Transcriptase according to the manufacturer’s instructions.

### Sequence-to-function mapping

Functional scores were quantified as the log2 of the fold-change enrichment of the variant reads-per-million (RPM) after the screen relative to its RPM in the virus library, i.e. log2 (Assay RPM/Virus RPM). Fit4Function models utilized the same design of the ML framework utilized for production fitness mapping (two-layer LSTM, custom early stopping, batch size of 500 variants, MSE error and Adam optimizer). Out of the 240K variants in the Fit4Function library, 90K were allocated for training and testing the ML function models (model construction) and 150K variants were held-out for validation of the MultiFunction approach. The training size for each function model was optimized independently. As with the production fitness model, the function models were assessed by correlation between the predicted and measured functional scores.

### MultiFunction library design

Using the previously generated fitness models of the production fitness and the five functional models described in the main text, we conducted an *in silico* screen of 10M randomly sampled 7-mer sequences to identify variants that are highly fit for all six traits. The threshold of high fitness for each function was arbitrarily set to the 50th percentile of each functional fitness distribution from the Fit4Function screening data. The percentiles were calculated on the detected variants of each functional assay from the 90K model construction data set. To reduce false positive predictions (variants predicted above the thresholds due to model errors), we increased the filtration thresholds slightly when applied to the predictions. For example, if the measured threshold is at a fitness score of 2.5, we considered variants predicted to have fitness >*2.5+shift*. The *shift* in applied thresholds is arbitrarily set to be 5% of the fitness dynamic range of each function. The thresholds were then used to filter out the 10M variants that were run through the six functional prediction models.

Out of the variants predicted to pass the six modified thresholds, we sampled 30K variants to be included in the MultiFunction library. The 30K variants were each represented by two 7-mer replicates. The MultiFunction library also included (1) a positive control set (3K) that was drawn from the subset of the 150K Fit4Function validation set that met the six conditions on the actual measurements (without modifying the thresholds), (2) a set of 10K variants randomly sampled from the Fit4Function 240K core variants as background controls representing the high production fitness space, (3) a set of 3K calibration variants present in the Fit4Function library (and the modeling library) to be used as background controls representing the entire (unbiased) sequence space, and (4) 1K stop codon containing sequences.

### MultiFunction library validation

The MultiFunction library was synthesized, virus was produced, and the five liver-related functions were screened in the same way the Fit4Function library was processed. We quantified the success rate of the MultiFunction library in terms of hit rate, i.e. out of the 30K variants predicted to meet the six criteria, what percentage satisfied the six criteria when the MultiFunction library was screened on those functions (predicted positive versus measured positive). To determine whether a variant meets specific functional criteria, we compared the distribution of that function for the MultiFunction variants against the positive control set. For a variant to be considered a hit for a specific function, its measured value should be above the mean-2SD (standard deviations) of the positive control set measured in the same experiment. A variant is considered a hit in calculating the MultiFunction hit rate only if it is a hit for all six functions; a variant that meets five or fewer conditions is not considered a hit.

The hit rate of the Fit4Function space is the number of non-control variants from the Fit4Function library measured to pass the six thresholds (without the prediction marginal shifts used for MultiFunction variant design) divided by the number of non-control variants in the library. The hit rate for the uniform sequence space can be estimated as the hit rate in the Fit4Function library (representing the high production fitness space – all the low production fitness variants were filtered out from the selection), relative to the percentage of the space occupied by the high production fitness variants. Uniform hit rate = Fit4Function hit rate × High production fitness ratio = 7.1% × 40.8% = 2.9%.

### Individual capsid characterization

Individual capsids were cloned into iCAP-AAV9 (K449R) backbone (GenScript), produced as described above, and administered to C57BL/6J (The Jackson Laboratory, 000664) mice at a dose of 1×10^10^ vg/mouse (*n*=5/group). Three weeks later, three separate lobes of the liver were collected for RNA extraction and a single lobe per mouse was fixed in 4% PFA.

For microscopy, fixed liver tissues were sectioned at 100 μm using a Leica VT1200 vibratome. Sections were mounted with ProLong™ Gold Antifade Mountant with DAPI (ThermoFisher, P36931). Liver images were collected using the optical sectioning module on a Keyence BZ-X800 with a Plan Apochromat 20X objective (Keyence, BZ-PA20). Three images were taken for each animal (n=5/group) and compared to a no injection control (*n*=3 animals). In CellProfiler, nuclei were segmented and DAPI^+^ nuclei were identified using a threshold on DAPI intensity determined from the no injection control. Each DAPI^+^ nuclei was then quantified with the median pixel intensity in the GFP channel.

For assessment of liver transduction by quantitative RT-PCR, total RNA was recovered using TRIzol (Invitrogen, 15596026) following the manufacturer’s instructions. Total RNA was then processed through an RNeasy kit (Qiagen, 74106) followed by on-column DNA digestion. RNA was converted to cDNA using Maxima H Minus Reverse Transcriptase (Thermo Scientific, EP0751) according to manufacturer instructions. Afterwards, qPCR was used to detect AAV encoded RNA transcripts with the following primer pair (5’-GCACAAGCTGGAGTACAACTA-3’) and (5’-TGTTGTGGCGGATCTTGAA-3’) and the following primer pair for GAPDH (5’-ACCACAGTCCATGCCATCAC-3’) and (5’-TCCACCACCCTGTTGCTGTA-3’).

THLE and HepG2 cells were seeded in a 96-well plate the day before adding the AAVs at 5000 vg/cell. For binding assays, viruses were diluted in media and incubated with cells at 4°C with gentle shaking for one hour. After incubation, cells were washed three times with PBS to remove unbound virus and treated with proteinase K to release viral genomes for qPCR quantification. For transduction assays, cells were incubated with the AAVs for 24 hours at 37°C and assayed with Britelite plus (Perkin Elmer, cat#6066766) following the manufacturer’s protocol.

## Data and Code availability

Data and code for algorithms described in the main text or Methods that are necessary for reproducing the study will be made available upon publication. Individual capsid plasmids described in this study will be made available through Addgene.

## Acknowledgements

We thank the members of the Deverman laboratory for continuous, extensive discussions of the project; Ray Jones, D. R. Mani and Andrew Barry for reviewing the manuscript; Sandrine Muller for discussions; and Andrew Steinsapir for assistance organizing the Fit4Function library macaque study.

## Author contributions

Conceptualization: BED, FEE

Methodology: BED, FEE, KYC, ATC, YAC Investigation: FEE, KYC, ATC, SP, IGT, QH, QZ, JJ, CK, PPB, BZ, MP, AFT Formal analysis: FEE, ATC

Visualization: ATC, FEE, YAC, BED

Funding acquisition: BED, FEE, KL, YAC, AFT

Supervision: BED, FEE

Writing – original draft: FEE, YAC, BED, ATC

Writing – review& editing: FEE, YAC, BED, ATC, AFT

## Supplementary Figures

**Supplementary Fig. 1.**
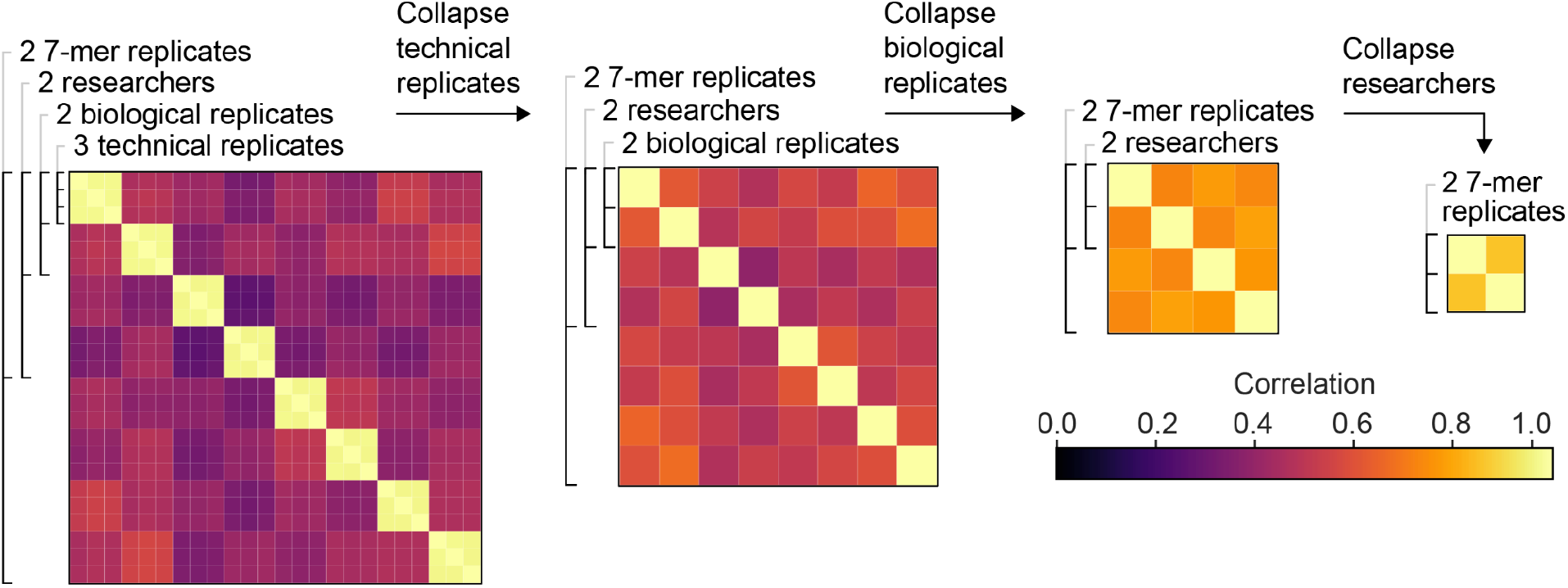
Production fitness replication quality improves upon hierarchical aggregation of replicates. Replication quality between replicates, where replication quality is defined as the Pearson correlation of log2 reads per million (RPM) between replicates. Data is aggregated (averaged) by technical replicates, then biological replicates, then by researchers, with replication quality increasing as replicates are collapsed.

**Supplementary Fig. 2.**
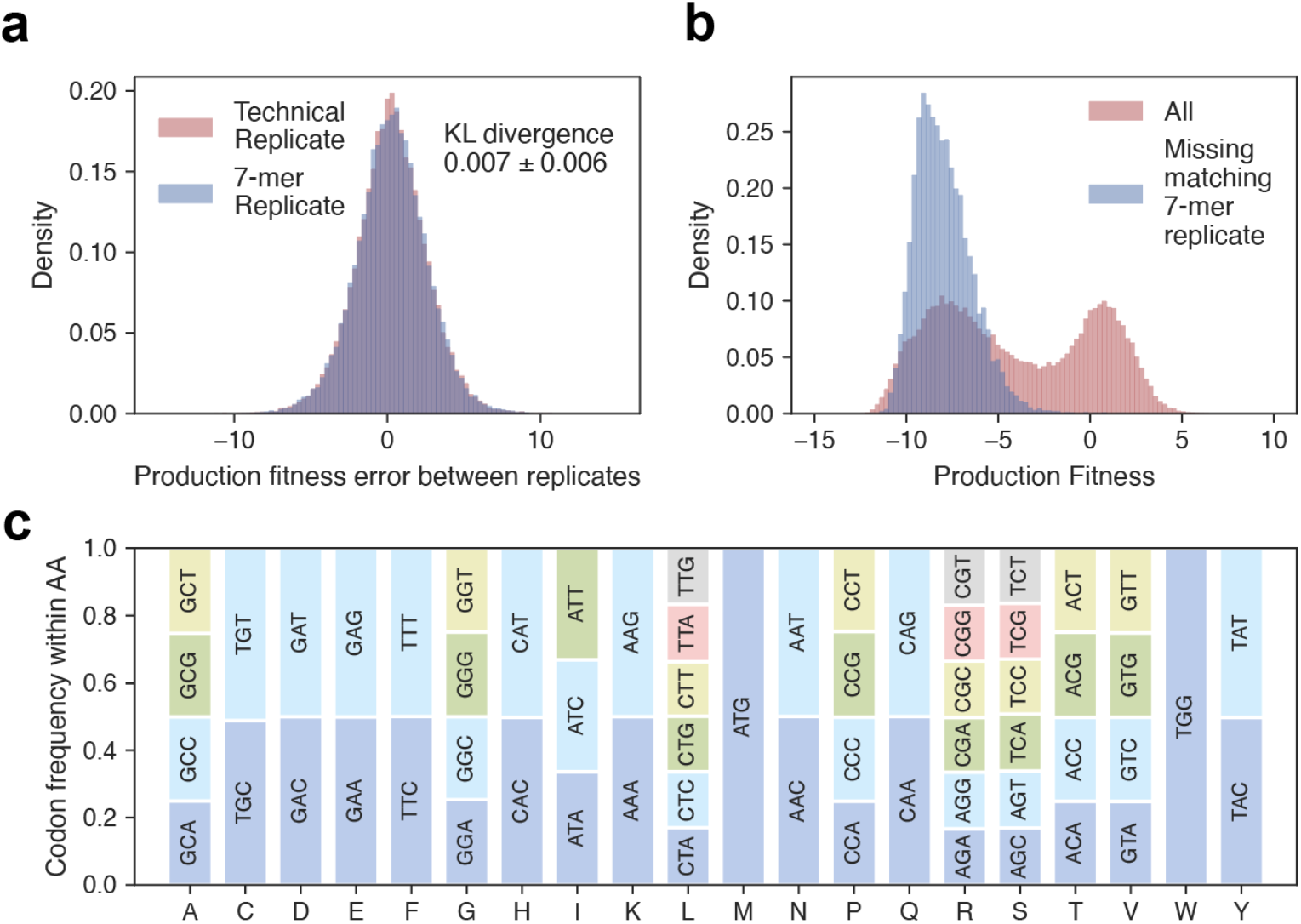
Codon usage of 7-mer insertions minimally affects capsid production fitness. The distribution of the difference in production fitness scores measured between 7-mer replicate pairs and between technical replicate pairs are similar (Kullback–Leibler divergence = 0.006±0.007). (**b**) The variants with a single 7-mer replicate detected (missing matching 7-mer) had production fitness scores on the low end of the production fitness bimodal distribution. Of the 13,217 7-mer replicates where only one of the two 7-mer sequences was detected in the virus (20.5% of the 64,500 amino acid variants), >99% had fitness scores on the low end of the fitness distribution, suggesting that the missing replicates were not detected due to low abundance. (**c**) Codon usage distribution in the modeling library follows the expected uniform distribution for each amino acid.

**Supplementary Fig. 3.**
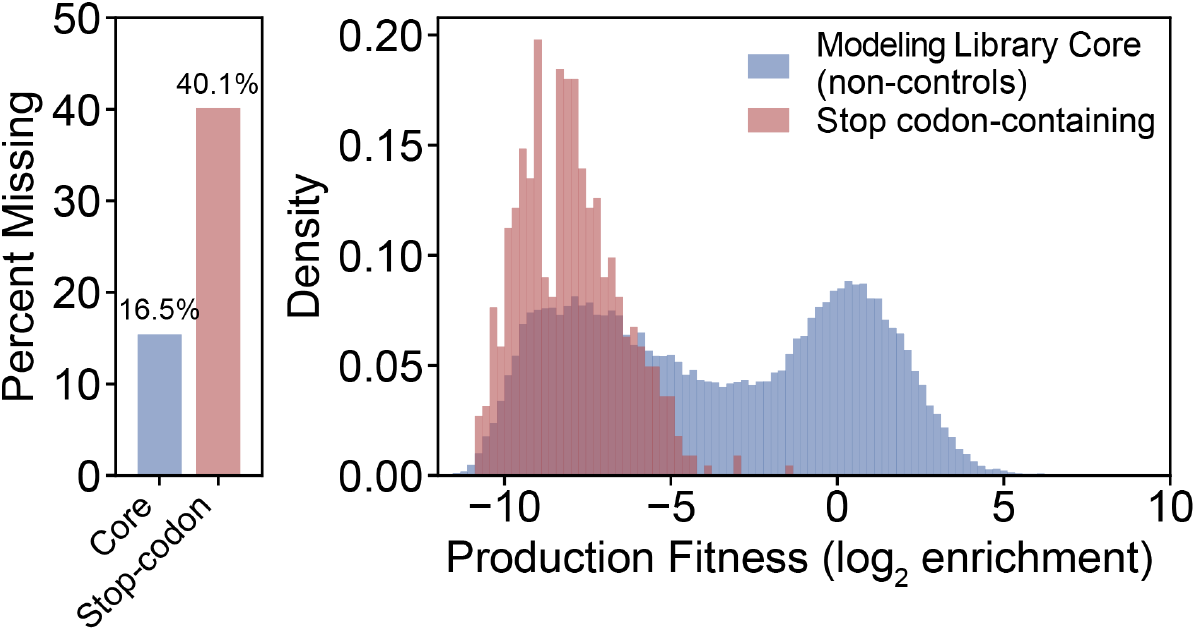
Distinguishing high and low production fitness distributions. The production fitness of detected stop-codon-containing variants in the modeling library, presumably arising due to cross-packaging, versus the production fitness landscape of the detected non-control variants (7-mer replicates not aggregated). Of the stop codon-containing sequences, 40.1% were not detected in the virus library.

**Supplementary Fig. 4.**
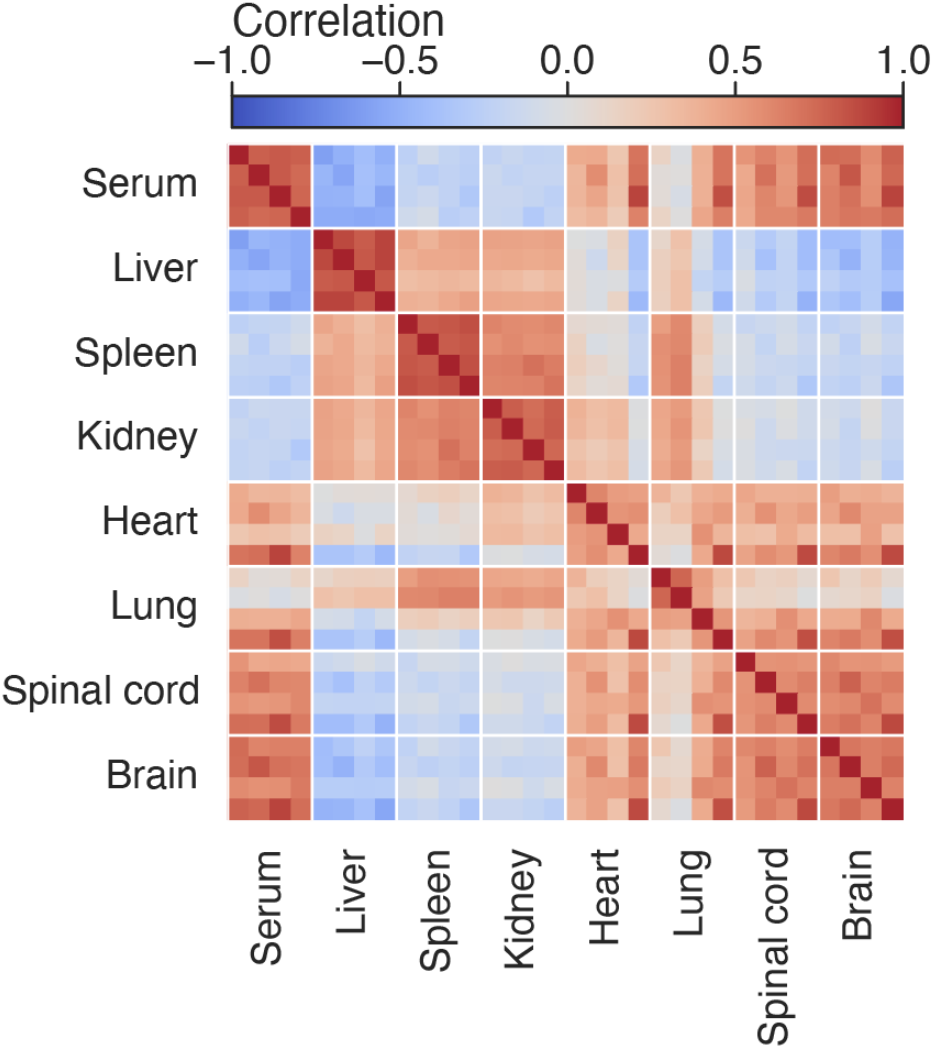
Fit4Function variant biodistribution correlation between organs. Pairwise Pearson correlation of organ biodistribution fitness (log2 enrichment scores) between C57BL/6J mice (*n* = 4 animals).

**Supplementary Fig. 5.**
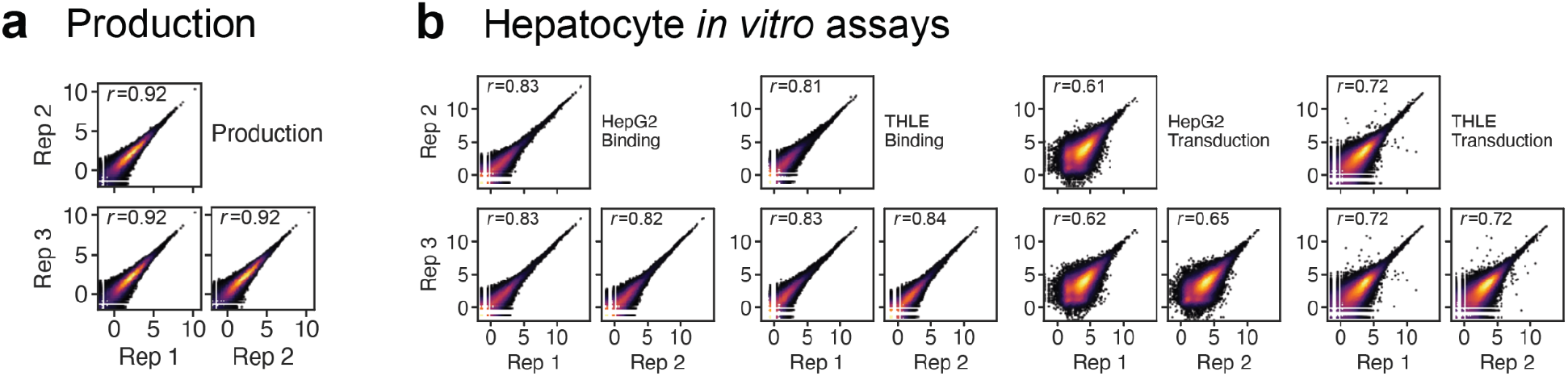
Replicability of the *in vitro* assays used to screen the Fit4Function library. Pairwise correlations between biological triplicates for (**a**) production fitness and (**b**) HepG2 binding or transduction and THLE binding or transduction.

**Supplementary Fig. 6.**
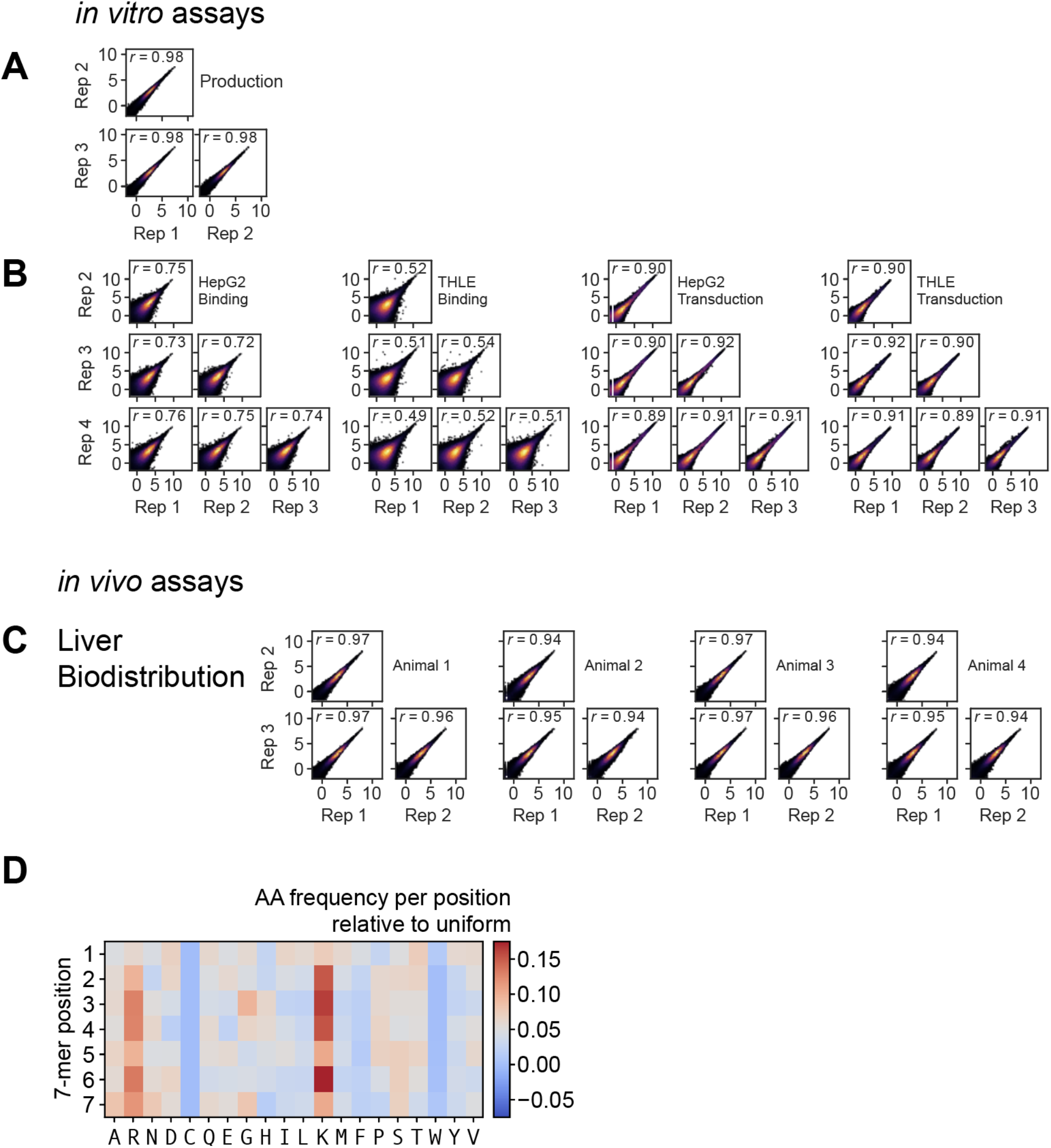
Replicability of the MultiFunction library across *in vitro* and *in vivo* assays. (**a**) Production fitness. (**b**) *In vitro* human cell binding and transduction. (**c**) *In vivo* liver biodistribution in C57BL/6J mice. (**d**) The amino acid distribution by position for the variants in the MultiFunction library.

**Supplementary Fig. 7.**
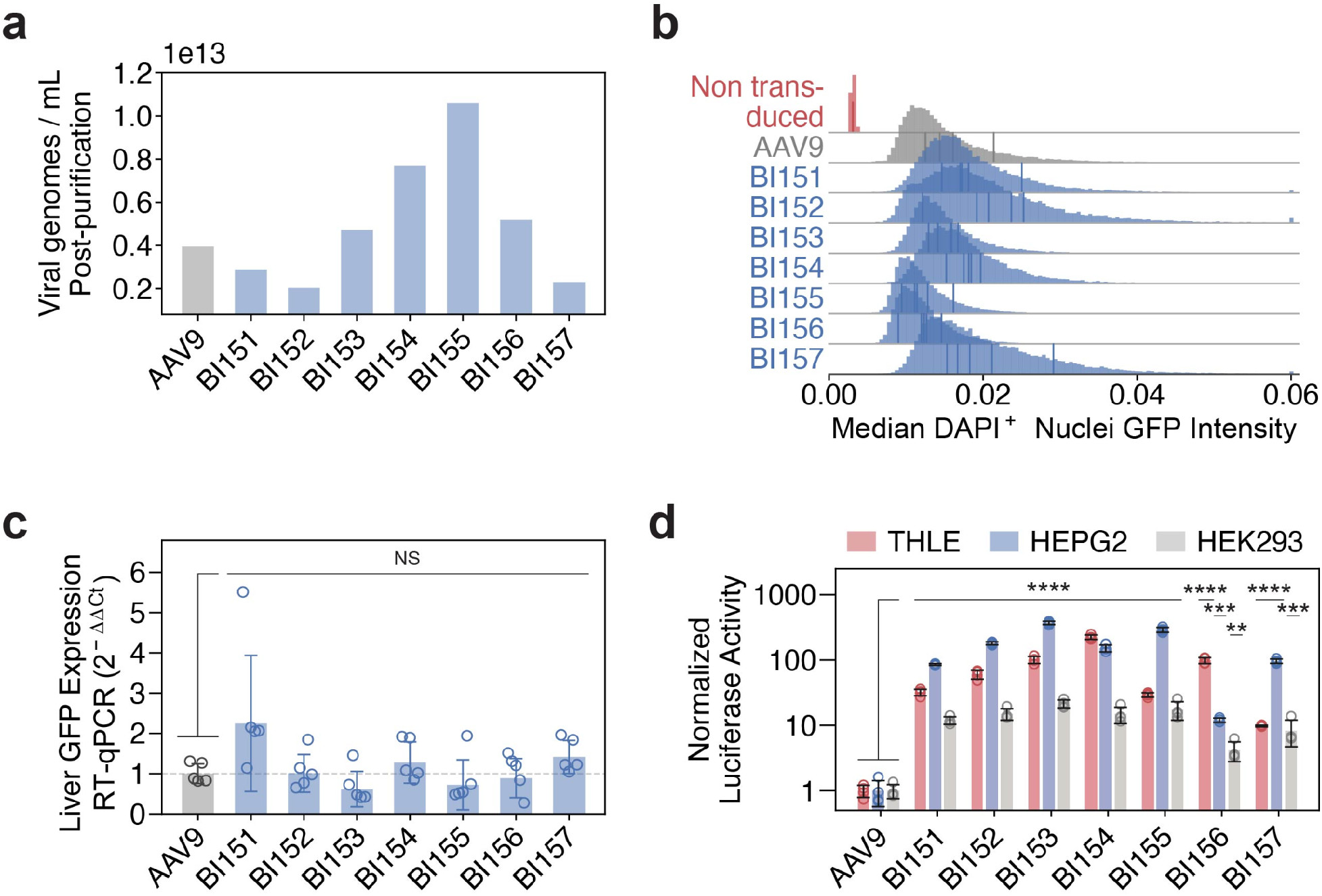
Individual assessment of liver MultiFunction capsids for production and cell transduction. (**a**) The production yields from 200 mL of cell culture for each of the individually manufactured capsids. (**b**) C57BL/6J liver transduction by AAV9 or MultiFunction capsids. Mice were injected with 1×10^10^ vg of the indicated capsid packaging AAV-CAG-GFP-2A-Luc-WPRE-pA and assessed for GFP expression three weeks later (*n* = 5 mice for each AAV treatment condition, *n* = 3 mice for the no AAV control, mean ± s.d., all BI capsids were not significantly different from AAV9 in unpaired, one-sided t-tests with Bonferroni correction). The distributions of median GFP pixel intensity per DAPI^+^ nuclei, combined across *n* = 5 animals for each AAV treatment condition, and *n* = 3 animals for the no AAV control are shown. The vertical lines within each distribution represent the mean of each animal. (**c**) AAV9 or the indicated capsid was used to transduce C57BL/6J mice at 1×10^10^ vg/mouse. Three weeks after AAV administration, liver transduction was measured by RT-qPCR of AAV transcripts from extracted tissue. △△Ct was obtained by normalizing against the reference gene (GAPDH), and then against the AAV9 control (*n* = 5 animals/group; mean ± s.d., unpaired, one-sided t-tests on log-transformed values, and Bonferroni corrected for multiple-hypotheses). (**d**) Human liver cell line (THLE, HepG2) and HEK293 transduction 24 hours after exposure to 5000 vg/cell of each indicated capsid packaging AAV-CAG-GFP-2A-Luc (*n* = 4 per group, mean ± s.d., \*\**p*<0.01, \*\*\**p*<0.001, \*\*\*\**p*<1e-4 unpaired one-sided t-tests on log-transformed values, and Bonferroni correction for multiple-hypotheses). Luciferase relative light units were normalized to AAV9.

**Supplementary Fig. 8.**
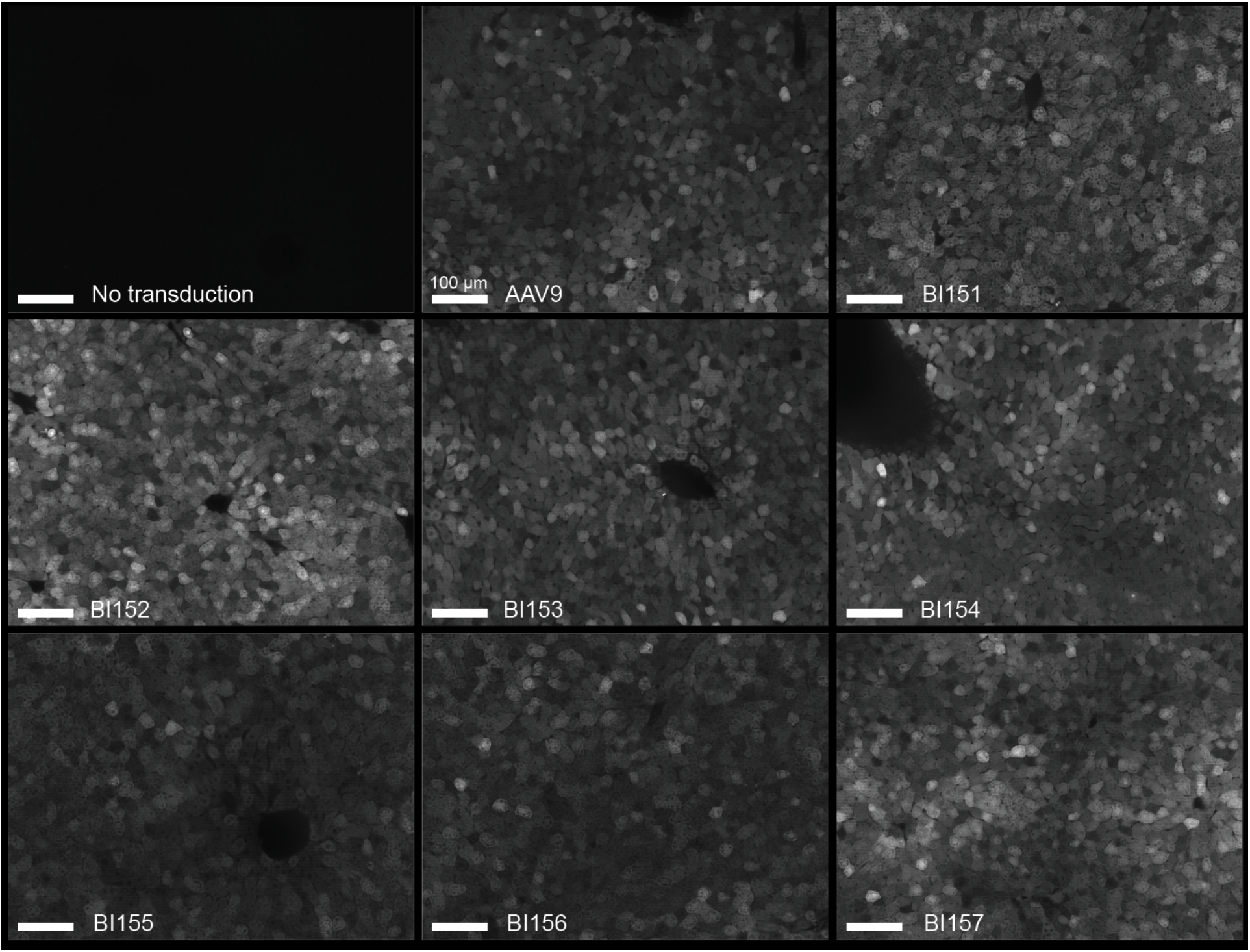
*In vivo* mouse liver transduction by each MultiFunction capsid. Representative GFP images of liver slices for the no AAV control, AAV9, and BI variants. Images were chosen from the median replicate of the median animal per condition. All images were taken at the same exposure and rescaled to the same intensity range. Scale bar in all images = 100 μm.

